# Integral Gene Drives: an “operating system” for population replacement

**DOI:** 10.1101/356998

**Authors:** Alexander Nash, Giulia Mignini Urdaneta, Andrea K. Beaghton, Astrid Hoermann, Philippos Aris Papathanos, George K. Christophides, Nikolai Windbichler

**Affiliations:** Department of Life Sciences, Imperial College London, Sir Alexander Fleming Building, South Kensington Campus, London, United Kingdom; Centre of Functional Genomics, Department of Experimental Medicine, University of Perugia, Perugia, Italy; Department of Entomology, Robert H. Smith Faculty of Agriculture, Food and Environment, Hebrew University of Jerusalem, Rehovot, Israel; to whom correspondence should be addressed (,)

## Abstract

First generation CRISPR-based gene drives have now been tested in the laboratory in a number of organisms including malaria vector mosquitoes. A number of challenges for their use in the area-wide genetic control of vector-borne disease have been identified. These include the development of target site resistance, their long-term efficacy in the field, their molecular complexity, and the practical and legal limitations for field testing of both gene drive and coupled anti-pathogen traits. To address these challenges, we have evaluated the concept of Integral Gene Drive (IGD) as an alternative paradigm for population replacement. IGDs incorporate a minimal set of molecular components, including both the drive and the anti-pathogen effector elements directly embedded within endogenous genes – an arrangement which we refer to as gene “hijacking”. This design would allow autonomous and non-autonomous IGD traits and strains to be generated, tested, optimized, regulated and imported independently. We performed quantitative modelling comparing IGDs with classical replacement drives and show that selection for the function of the hijacked host gene can significantly reduce the establishment of resistant alleles in the population while hedging drive over multiple genomic loci prolongs the duration of transmission blockage in the face of pre-existing target-site variation. IGD thus has the potential to yield more durable and flexible population replacement traits.

## Introduction

Homing gene drives were first proposed 15 years ago as potential tools for enabling the genetic engineering of natural populations (Burt 2003). After showing promise in a number of proof-of-principle studies (Chan et al. 2011; Windbichler et al. 2011; Chan et al. 2013; Simoni et al. 2014; V. M. Gantz and Bier 2015), first implementations highlighting their potential for the use to control disease vectors by population suppression and replacement were recently demonstrated in two species of malaria vector mosquitoes (A. Hammond et al. 2016a; Valentino M. Gantz et al. 2015a). Gene drive research is currently focused on two main areas: (i) studying the nature of target site resistance (A. M. Hammond et al. 2017a; Champer et al. 2017b; KaramiNejadRanjbar et al. 2018) to mitigate its eventual rise, and, conversely, (ii) reducing or counteracting the invasive potential of gene drives, in order to minimize the perceived or actual risk associated with the technology. The former strand of research is centered on improving regulatory elements to contain/confine nuclease activity to homing-relevant cell types, to identify target sites that are intolerant to drive-inactivating mutations and on the addition of further components to the drive constructs such as multiple gRNAs (Noble et al. 2017; Marshall et al. 2017; Champer et al. 2018a) or factors that bias repair towards the desired homology-directed pathway. For limiting gene invasiveness, a number of schemes have been proposed, for example linking multiple driving and non-driving CRISPR/Cas9 transgenes into a chain where the spread of each construct depends on the allele frequency of the prior link in the chain (Noble et al. 2016; Esvelt and Gemmell 2017).

Although gene drive research is now mainly centered on drives built using the CRISPR genome editing toolset, proposed strategies adhere to the classic transgene paradigm, namely the use of pre-characterized promoter and terminator elements, each driving tissue-specific transgene expression as required for different functions in the germline (drive) and various somatic tissues (anti-pathogen effect), and with the resulting constructs inserted at arbitrarily chosen genomic sites, e.g. genes that are presumed to be neutral (Figure 1A). It has been observed that the resulting complex and large constructs can show unexpected behaviors (Valentino M. Gantz et al. 2015b; A. Hammond et al. 2016b), possibly resulting from the interaction between engineered components with each other, their non-native genomic context, and the fact that isolated regulatory elements may not fully recapitulate expression patterns of the endogenous loci they were derived from. Moreover, while a number of modelling studies have mapped out the ideal characteristics and the resulting predicted theoretical behavior of gene drives, what is often neglected are the practical implications and limitations for the construction of gene drives based on those schemes. For example, each and every one of the molecular components (promoters, gRNAs, fluorescent markers etc.) of complex constructs such as those that carry multiple anti-pathogen effectors or those designed around the use of multi-gRNA arrays have all to satisfy regulatory requirements. Along the same lines, limiting the propagation of gene drives by inserting them into repetitive genomic regions while attractive in principle presents a formidable genome engineering challenge (Min et al. 2017).

**Figure 1.**
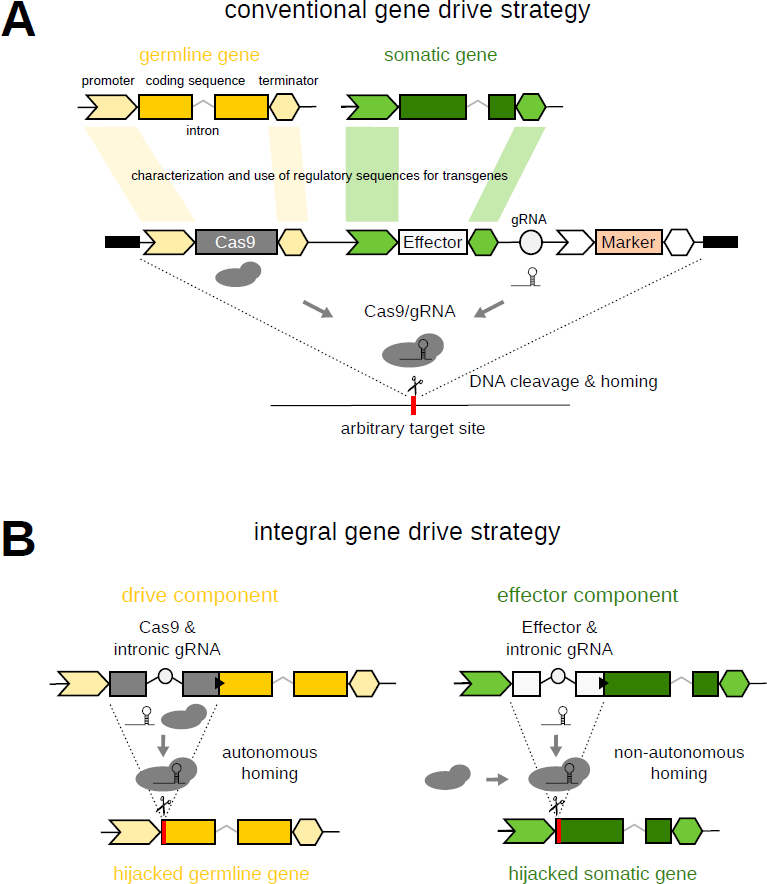
Schematic overview of the molecular constructs enabling both conventional (A) and integral (B) gene drive approaches for population replacement. The black triangle indicates the 2A ribosomal skipping signal and the gRNA locus is indicated as a circle.

Blocking parasite development in genetically modified mosquito vectors is an area intensively researched long before efficient gene drive had first been demonstrated (Ito et al. 2002a). The modification of vector genes involved in immunity and vector-parasite interactions or the introduction of exogenous effectors such as antimicrobial peptides and antibodies specifically targeting the parasite are the two cardinal approaches to interfere with *Plasmodium* transmission. A growing set of anti-pathogen effectors now exist (Kim et al. 2004; Ito et al. 2002b; Corby-Harris et al. 2010; Isaacs et al. 2011; Jasinskiene et al. 2007), yet these traits have been assessed exclusively against laboratory strains of *Plasmodium falciparum* or the rodent parasite *Plasmodium berghei*. Thus, the efficacy of these effectors against genetically diverse polymorphic isolates of the parasite is currently unknown. The necessary experiments can realistically only be performed in a disease endemic setting as they require the recruitment of gametocyte carriers from the human population. However, population-replacement strains as currently envisioned carry one or multiple anti-pathogen effector traits directly coupled to the endonuclease, and hence cannot be tested in the absence of gene drive complicating this crucial step (Figure 2A). Alternatively, standard transgenic effector strains must first be generated and used to perform these pilot experiments, which would require further genetic engineering steps (possibly altering their properties) to enable gene drive later on. This also highlights the fact that, despite ongoing discussions (Z. Adelman et al. 2017), no clear pathway for safely testing gene drives has emerged that would allow research to progress step by step from lab to field deployment. It was not until 2017 that the first transgenic mosquito strain, carrying a dominant male-sterilizing transgene (Windbichler, Papathanos, and Crisanti 2008), was imported by the Target Malaria consortium to an African partner nation, after completing a regulatory pathway that lasted nearly 2 years. Indeed, it is expected that gene drive strains will face a significantly tougher and prolonged regulatory pathway compared to strains harboring such a conservative genetic modification.

**Figure 2.**
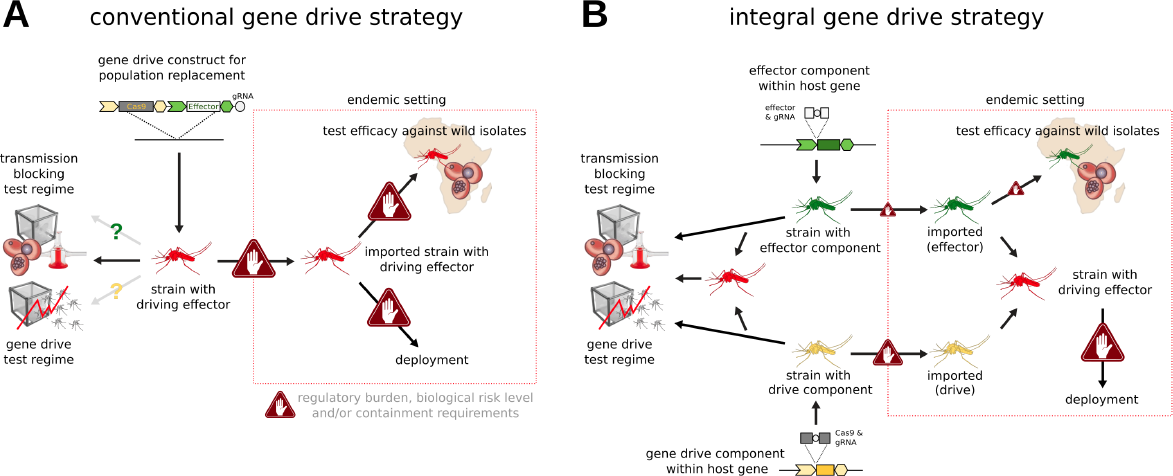
A comparison of the steps and requirements for lab and field testing of both conventional (A) and integral (B) gene drive strategies for population replacement of malaria vectors.

These foreseeable regulatory and operational challenges must therefore inform the design of gene drives, including minimizing their molecular complexity and allowing each of their components to be assessed independently. Such an approach could accelerate the pace of gene drive research and testing as it moves closer to the field. Here we evaluate a novel strategy, Integral Gene Drive (IGD), for implementing population replacement that is summarized in Figures 1B and 2B. IGD is specifically conceived with the aforementioned molecular, population dynamic, practical, operational and regulatory challenges in mind.

### IGD drive components

In contrast to the design of conventional population replacement constructs that reflects the classic transgene paradigm (Figure 1A), IGDs integrate the endonuclease coding sequence (e.g. Cas9) directly within an endogenous gene whose function and expression is confined to the male and/or the female germline where homing occurs (Figure 1B). The presence of Cas9 should ideally have no significant negative effect on the expression of the hijacked host gene. To guarantee accurate translation of both Cas9 and the endogenous host gene, their open reading frames are linked via the 2A ribosome-skipping signal, resulting in the production of two distinct functional polypeptide chains from a compound transcript. A similar strategy has been successfully used to generate endogenously driven reporter genes (Rojas-Fernandez et al. 2015). Both an N-terminal as well as a C-terminal fusion of Cas9 to the host gene is possible. However, one consequence of the former arrangement is that incomplete homing events or frameshift mutations inactivating Cas9 would also lead to the loss of the function of the hijacked host gene. Therefore, selection would be expected to maintain the integrity of the Cas9 open reading frame to a certain degree.

The regulatory elements of a number of germline genes have so far been validated in flies and mosquitoes. These include promoter elements derived from the vasa (Papathanos et al. 2009), nanos (Calvo et al. 2005), or beta2-tubulin (Catteruccia, Benton, and Crisanti 2005) genes and other loci (Akbari et al. 2014). These genes are thus ideal candidates for serving as hijacked host genes for the drive component. As shown in Figure 1B, the gRNA expression cassette can be located within an intron located inside the Cas9 coding sequence. Intronic gRNAs have been demonstrated previously (Ding et al. 2018; Kiani et al. 2014) and we recently explored this concept in insects (Nash et al, in preparation). Intronic gRNAs can either be promoterless and thus mirror the expression pattern of the hijacked host gene, or they can have their own RNA Polymerase III promoter element for ubiquitous expression (e.g. in cases the host gene is not itself expressed in the germline as in the case of the effector component described below). In all cases, targeted cleavage mediated by the gRNA and Cas9, which associate with each other in the germline, triggers homing. The drive component is thus an autonomously homing allele of an endogenous gene and is designed to spread in a population (at the expense of wild-type alleles of the same gene) with no other intended effect than seeding it with and increasing the allele frequency of the coupled Cas9 trait.

### IGD effector components

For simplicity, we consider here the effector to be an exogenous polypeptide that when expressed in the target tissue reduces or abolishes parasite development in the mosquito. Various mosquito tissues such as the midgut, the hemocytes and the salivary glands are at the interface of the vector-parasite interactions and are thus ideal candidates for hosting the expression of effectors. The fat body is another good candidate tissue, as its secretions into the hemocoel can directly interact with (Ito et al. 2002c) parasite oocyst and sporozoite stages. Analogous to the drive component, each effector is incorporated into an endogenous gene expressed in any of the above target tissues (Figure 1B) thereby hijacking that gene for the use of its regulatory regions. In mosquitoes, a limited number of regulatory regions driving transgene expression in these target tissues have been characterized, including the carboxypeptidase promoter shown to drive transgene expression in the midgut following a blood-meal and the vitellogenin promoter driving blood-meal induced expression in the fat body. However, genome-wide expression analyses have identified numerous additional genes specifically expressed in these tissues (Giraldo-Calderón et al. 2015). Indeed, IGD sidesteps the laborious process of first experimentally testing the temporal and spatial specificity of isolated promoter sequences and instead directly utilizes any such suitable loci for targeted insertion of effector transgenes. Again, an N-terminal fusion of the effector transgene with respect to the host gene can guard against incomplete homing events or frameshift mutations that lead to the loss of the function of the host gene, maintaining the integrity of the effector. Other approaches for achieving hijacking than the use of the 2A signal are also conceivable but are not discussed here in detail (Supplementary Figure 1). Each IGD effector transgene also carries a U6-driven intronic gRNA for triggering homing in the germline. Unlike the drive component however, the effector component does not encode an endonuclease and thus is not able to initiate gene drive on its own. Indeed, targeted cleavage mediated by the gRNA and Cas9 complex can only occur when the latter is provided in trans by the drive component. Cleavage in the germline triggers homing of the effector component, which is a non-autonomously homing allele of an endogenous gene and is designed to spread in a population with the intended effect of increasing the allele frequency of the coupled effector trait.

### Developing IGD traits

Drive and effector traits inserted at various suitable loci throughout the genome could be generated and tested independently from each other, including by different research teams (Figure 2B). On the one hand, the drive component can be specifically optimized in the laboratory for its propensity to induce homing, the faithfulness of Cas9 expression and to minimize the rate at which cleavage triggers the undesirable NHEJ or MHMR repair pathways. In addition, one would seek to reduce the fitness cost incurred by the expression of Cas9 itself or by its integration interfering with the function of the hijacked host gene. On the other hand, the effector component can be optimized in the laboratory for its efficacy in reducing or blocking parasite/pathogen development and transmission, its own intrinsic fitness cost and any negative effect it may have on the expression of the hijacked host gene. In addition, the rates of non-autonomous homing of the effector component, assisted by the presence of the IGD drive component or another source of Cas9, can be measured under laboratory containment. Laboratory crossing of the transgenic strains harboring the IGD drive and effector components would allow assessing the likely performance of these trait combinations in the field, including the spread of each modified allele and their resulting frequency in cage populations.

Field testing of the effector component would be aided by the simplicity of the genetic modification, i.e. insertion of a single effector coding sequence containing the intronic gRNA, with all other functions provided by the hijacked host gene. Given the inability of the effector component to spread autonomously, the regulatory threshold for such strains is expected to be lower than conventional fully-driving transgenes harboring a population replacement payload (Figure 2A). This would facilitate import of these strains to disease endemic countries and the swift testing of effector traits against polymorphic isolates of the parasite as well as within varying mosquito genetic backgrounds (Figure 2B). Strains harboring the IGD drive component are also conceived to be molecularly simple, although they will would be more difficult to import, regulate and deploy as they carry autonomous gene drive elements. However, the drive component on its own is not designed to have any phenotypic effect on fitness or vectorial capacity of the mosquito. Thus, an inadvertent release and spread of such a strain would not be expected to have any relevant effect on mosquito biology and carries a relatively lower risk.

The scientific working group on a pathway to deployment of gene drive mosquitoes for elimination of Malaria in Sub-Saharan Africa (James et al. 2018) recommends the use of fluorescent markers to track gene drive constructs pre and post-deployment. However, gene drives can decouple from genetically-linked fluorescent marker genes within a single generation of homing, potentially giving rise to type II errors during monitoring, arguably the most crucial error as it would suggest the absence of constructs in populations or regions in which active gene drives are in-fact propagating. Fluorescent markers, other than those used for transgenesis and that can be subsequently removed, are therefore to be avoided in our design and modelling of IGD population replacement, as they also increase molecular and regulatory complexity. We assume that molecular genotyping will be the only viable approach for gene drive monitoring. Replacement drives including IGDs, unlike suppressive drives, can be constituted as true-breeding strains which should facilitate the exclusive use of molecular markers during implementation.

### Exploring IGD population dynamics

To predict the behavior of one or multiple interacting IGD traits on the population level, we used a discrete-generation (non-overlapping) model comparing the dynamics of a classic replacement drive (Beaghton et al., 2017) to the dynamics of IGD, analyzing protection levels and allelic dynamics over time. To facilitate comparison to the conventional replacement drive model, we initially constructed a two-locus model with one drive component hijacking a germline gene (nuclease, locus 1) and one effector component hijacking a somatic gene (effector, locus 2). We then extended this to a three-locus two-effector model with a single gene drive component at locus 1 and effector components at two independent loci 2 & 3, assuming for the sake of simplicity that the effectors at the two different host genes have the same molecular biology and fitness parameters. As a baseline, we assume that if an individual has at least one effector component at locus 2 or 3, we consider it to be refractory against malaria. We evaluate the effectiveness of these different drive architectures by calculating the duration of protection, which is affected by the probabilities of different molecular processes such as homing and the formation or pre-existence of resistant alleles, by the fitness costs of the nuclease and the effector components, and by the efficacy of the effector. Protection is defined as the reduction in vectorial capacity, given by the sum of the genotype frequencies with at least one effector component at either locus times their degree of reduction on vector competence. A baseline set of parameter values (Table 1) was chosen to be consistent with existing published work on mosquitoes (Hammond et al. 2016) and for ease of comparison to the classical replacement drive model (Table 2).

**Table 1.** **Parameters and Baseline Values for the IGD Model.** An exemplar set of parameter values that is consistent with most of the extensive published work on mosquitoes (Hammond et al. 2016) has been chosen for both the classical gene drive model (Beaghton et al. 2017) and the IGD drive model. These parameters define various aspects of molecular biology, fitness and vector competence effects, and initial genotype frequencies. Parameters shared by both models are given the same baseline values to facilitate comparison (e.g., drive transmission, loss-of-function mutations during HDR, costs for nuclease and effector expression).

| Symbol | Parameter<sup>+</sup> | Baseline Value |
| --- | --- | --- |
| $cc_0$ | Initial release frequency of homozygous drive and effector construct | $10^{-4}$ |
| $dd_0$ | Initial release frequency of homozygous drive construct | 0 |
| $ee_0$ | Initial release frequency of homozygous effector construct | 0 |
| $rr1_0$ | Initial frequency of resistance (locus 1) | 0 |
| $rr2_0$ | Initial frequency of resistance (locus 2) | 0 |
| $rr3_0$ | Initial frequency of resistance (locus 3) | 0 |
| $d_1$ | Transmission rate of drive (locus 1) | 0.99 |
| $d_{2a}$ | Transmission rate of drive (locus 2) if drive component present in heterozygosity | 0.99 |
| $d_{2b}$ | Transmission rate of drive (locus 2) if drive component present in homozygosity | 0.99 |
| $d_{3a}$ | Transmission rate of drive (locus 3) if drive component present in heterozygosity (I) | 0.99 |
| $d_{3b}$ | Transmission rate of drive (locus 3) if drive component present in homozygosity (I) | 0.99 |
| $l_1$ | Resistance arising from HR at locus 1 | $10^{-4} \times (1/3)$ |
| $l_2$ | Resistance arising from HR at locus 2 | $10^{-4} \times (1/3)$ |
| $l_3$ | Resistance arising from HR at locus 3 (I) | $10^{-4} \times (1/3)$ |
| $u_1$ | Resistance arising from NHEJ at locus 1 | $0.5 \times (1/3)$ |
| $u_2$ | Resistance arising from NHEJ at locus 2 | $0.5 \times (1/3)$ |
| $u_3$ | Resistance arising from NHEJ at locus 3 (I) | $0.5 \times (1/3)$ |
| $L_1$ | Loss of gene function arising from HR at locus 1 | $10^{-4} \times (2/3)$ |
| $L_2$ | Loss of gene function arising from HR at locus 2 | $10^{-4} \times (2/3)$ |
| $L_3$ | Loss of gene function arising from HR at locus 3 (I) | $10^{-4} \times (2/3)$ |
| $U_1$ | Loss of gene function arising from NHEJ at locus 1 | $0.5 \times (2/3)$ |
| $U_2$ | Loss of gene function arising from NHEJ at locus 2 | $0.5 \times (2/3)$ |
| $U_3$ | Loss of gene function arising from NHEJ at locus 3 (I) | $0.5 \times (2/3)$ |
| $s_m$ | Cost of loss of gene function | 1 |
| $s_{d1}$ | Cost of hijacking target locus 1 (Drive) | 0 |
| $s_{d2}$ | Cost of hijacking target locus 2 (Effector) | 0 |
| $s_{d3}$ | Cost of hijacking target locus 3 (Effector) (I) | 0 |
| $s_n$ | Cost of expressing nuclease at locus 1 | 0.05 |
| $s_{e2}$ | Cost of expressing effector at locus 2 | 0.1 |
| $s_{e3}$ | Cost of expressing effector at locus 3 (I) | 0.1 |
| $h_m$ | Dominance coef. for loss of gene function | 0.2 |
| $h_{d1}$ | Dominance coef. for hijacking at locus 1 | 0.5 |
| $h_{d2}$ | Dominance coef. for hijacking at locus 2 | 0.5 |
| $h_{d3}$ | Dominance coef. for hijacking at locus 3 (I) | 0.5 |
| $h_n$ | Dominance coef. for expressing nuclease | 0.5 |
| $h_{e2}$ | Dominance coef. for expressing effector (locus 2) | 0.5 |
| $h_{e3}$ | Dominance coef. for expressing effector (locus 3) (I) | 0.5 |
| $h_{rc1}$ | Dominance coef. for refractoriness (1 effector allele) | 1 |
| $h_{rc2}$ | Dominance coef. for refractoriness (2 effector alleles) (I) | 1 |
| $h_{rc3}$ | Dominance coef. for refractoriness (3 effector alleles) (I) | 1 |
| $r_c$ | Homozygous degree of refractoriness | 1 |

<sup>+</sup> I – Three locus model only

**Table 2.**
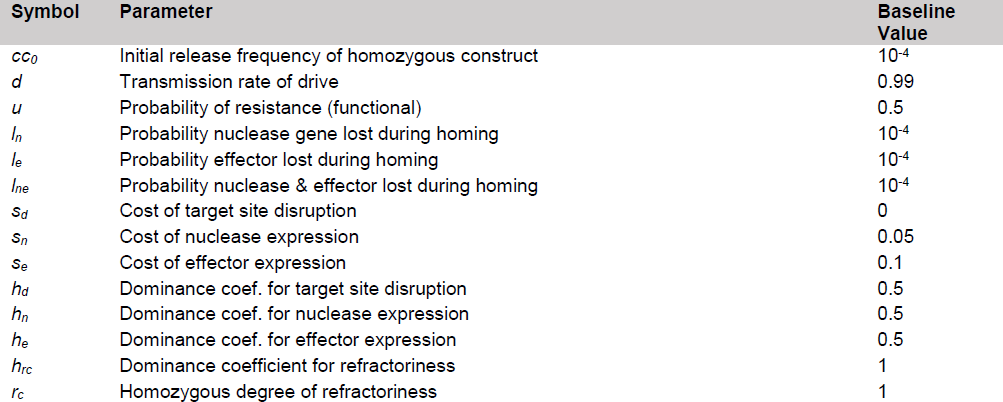
**List of Parameters and Baseline Values for ‘Classical’ Model.** An exemplar set of parameter values, consistent with published work conducted in relation to classical replacement gene drives (Beaghton et al. 2017c).

A comparison of a conventional replacement strategy and the IGD two-loci model is summarized in Figure 3. We find that, using identical baseline values (Tables 1 & 2) to facilitate comparison (e.g., drive transmission, loss-of-function mutations during HDR, costs for nuclease and effector expression), the IGD strategy conveys 95% protection for 81 generations, compared to 39 generations for the classical replacement drive. This translates to approximately 4.5 years of protection against malaria transmission, while protection given by a classical gene drive lasts approximately 2.1 years (Depinay et al. 2004; Mordecai et al. 2013). As in the conventional strategy, resistant alleles are generated at each locus, eventually replacing the constructs since they do not carry the cost associated with expressing either the nuclease (locus 1) or the effector (locus 2). If costs of expression due to the nuclease (*s_n_*=0.05) are less than for the effector (*s_e2_*=0.1) resistance replaces the transgene faster at the effector locus, while the nuclease persists for longer in the population, whereas in the conventional strategy the compound cost for expressing both causes the construct to be lost rapidly. We find that at the same rate of formation of resistant alleles, their impact is reduced in the IGD strategy since mutations that result in a loss of function of the hijacked endogenous target gene and are selected against – unlike the conventional strategy which assumes the target to be neutral.

**Figure 3.**
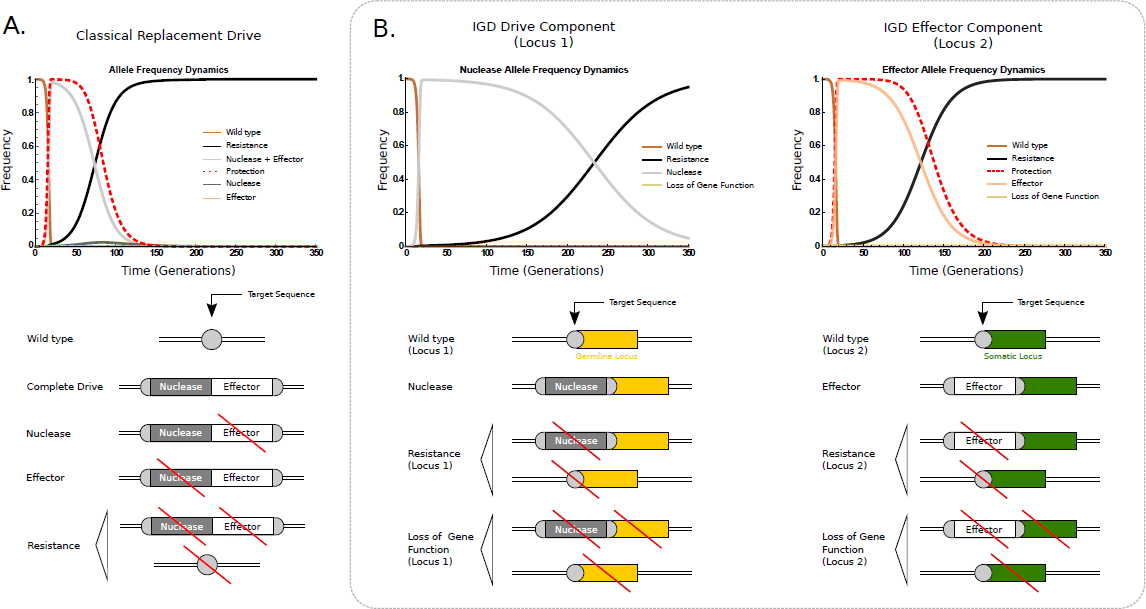
Comparison of allele frequency dynamics of the conventional (A) and integral (B) gene drive strategies using baseline parameter values. Integral drive dynamics is displayed here in terms of the modelled behavior of the constituent components at the two loci. The dashed red line shows the proportionate reduction in vectorial capacity of the target population.

In order to determine which parameters have the strongest effect on the duration of protection, we varied each while retaining all others at baseline values (Supplementary Figure 2). We find similar dependencies as for the classical model, however for a number of parameters IGD appears more robust with minor or no effects on the duration of protection evident. For example, increasing the proportion of resistance and loss of function alleles formed by either NHEJ or aberrant HDR at locus 1 within a biologically sensible range does not affect the duration of protection. This can be explained by the main effect the drive component (locus 1) has on locus 2, which is to convert wild-type alleles to effector alleles in heterozygous individuals allowing the effector to rapidly propagate in the population and to establish protection before resistance at locus 1 takes over. Protection starts decreasing only when resistant alleles start forming at locus 2 and eventually replace effector alleles in the population. The eventual subsequent loss of the drive allele at locus 1 and its replacement by resistant alleles is no longer of any consequence to the duration of protection because the conversion of wild-type alleles at locus 2 into effector has already taken place.

We found that the existence of pre-existing resistant alleles at the effector locus among the factors that most significantly reduce the duration of protection (Supplementary Figure 2). By contrast, levels of protection begin to crash only when initial resistance at locus 1 approaches 80%, i.e. that target sequence represents a minor allele. Should pre-existing resistance alleles occur with a frequency of 10% at both loci, 95% protection is reduced from 81 generations to 15 generations. Pre-existing resistance must be assumed to be present in significant proportions in many target species including mosquitoes (Anopheles gambiae 1000 Genomes Consortium et al. 2017), and modelling has already shown that resistant alleles arising from standing genetic variation are generally more likely to contribute to resistance than from new mutations induced by the drive (Unckless et al 2017). Having investigated the effect of pre-existing resistance alleles in a two-locus IGD model, which showed that only the effector locus is particularly sensitive to pre-existing resistance, we considered next our three-locus two-effector model. Deploying two-effector or multi-effector strains should sustain protection for longer, since for the protection to disappear, resistance will need to develop or pre-exist for the effector at both loci in a significant fraction of the population. We find that releasing a two-effector driving strain into a population without pre-existing resistance yields extended protection of 103 generations. With pre-existing resistant alleles (10% allele frequency at all 3 loci), 95% protection lasts for 38 generations (Figure 4), a significant increase in the duration of protection when compared to a single-effector strain release under identical conditions.

**Figure 4.**
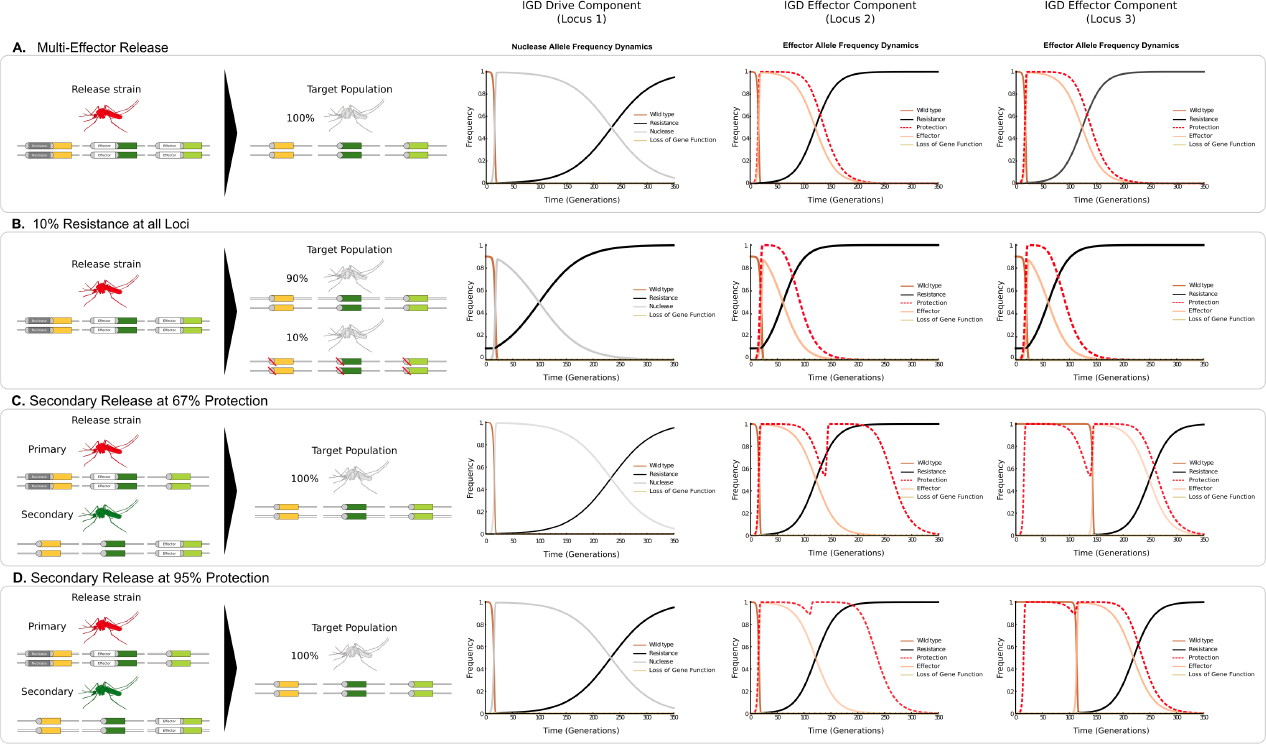
Dynamics of the 3-locus 2-effector model assuming the release of transgenics for all loci (locus 1: drive, locus 2 & 3: effectors) into either a population that is wild-type for all loci (A) or that carries pre-existing resistance at all loci at 10% allele frequency (B). Staggered release of the second effector strain at 95% (C) and 67% (D) protection, whilst all other parameters are maintained at their respective baseline value. Integral drive dynamics are displayed here in terms of the modelled behavior of the constituentcomponents at the 3 loci. The dashed red line shows the proportionate reduction in vectorial capacity of the target population.

A second effector component allele at a separate locus could also be introduced after a given time to extend the duration of protection. The second effector is driven through the population when in the presence of the nuclease allele at locus 1, extending the duration of protection until the time when the second effector is in its turn replaced by resistant alleles. Figure 4 shows the boost in duration of protection for these baseline parameters (*h*_n_=0.05, *h*_e2_=*h*_e3_=0.1) when the second effector is added at the time when the level of protection from the first effector has dropped to 67% and to 95%. When protection drops below a certain level, effector release restores maximal protection. The lower the cost of expression of the nuclease (*s_n_*), the longer it will be present in the population and the longer protection can be extended. The condition for full protection to be achieved after release of the second effector at locus 3 is that the allelic frequency of the nuclease is greater than ≅55% at the time of release. If the second effector is released before protection starts significantly decreasing, it can optimally prolong the duration of maximal protection. We find that the ideal time for secondary effector release at locus 3 is when protection drops below 95%.

### Testing and deployment of IGDs

The scientific working group on gene drives recommends a stepwise pathway for the deployment of gene drive mosquitoes i.e. to progress testing from laboratory studies, possibly involving large indoor and outdoor cage trials, to small scale isolated and open releases to full scale open releases (James et al. 2018). However, it is not always clear how limiting drive propagation can be guaranteed with conventional gene drive designs, as the level of ecological or geographical isolation achievable at different release sites is yet to be fully understood. The modularity and interdependence of IGD components does provide a straightforward pathway for moving testing from self-limited to self-sustaining traits in the field by modulating the propensity to spread in the population (Figure 5). First, an inundative release of an effector strain alone would allow to assay (by recapture) mosquito fitness and performance under field conditions and to detect any unintended effects prior to deployment. When tested in the absence of a drive component, effector strains will not convert the field population, permitting safe testing of individual effector components. A second scenario consists of releasing a limited-drive strain, containing an effector component and a non-driving source of Cas9, which can trigger a limited and local spread of the effector trait and allow evaluation of its drive performance and perhaps its epidemiological effect in reducing disease transmission. This Cas9 source permits super-Mendelian inheritance of effector components within the field population but is itself inherited at a Mendelian rate, and modelling suggests that both would be lost. This strategy therefore facilitates the testing of effector component homing in the field, without the perceived risk posed by using a driving source of Cas9. Modulating the allele frequency of the non-driving Cas9 trait via inundative releases of varying magnitudes would allow to control the expected level of spread and the resulting allele frequency of the effector. The effect of these two first strategies is self-limiting, perhaps allowing a test site to be re-used following the dissipation of the released alleles. When individual traits have been sufficiently tested separately and in conjunction in the laboratory, as well as in self-limiting pilot experiments in the field, one can consider the release of the fully driving strain carrying both drive and effector component alleles. The release of such a transgenic strain would then trigger full population-wide gene drive in the field and propagation according to the previously described dynamics. It is important to highlight that, unlike conventional designs, here the performance and behavior of the IGD effector trait is likely to be unchanged by the addition of the drive component. A regular population replacement strategy would require, for various stages of testing, different driving and non-driving constructs to be made that could display significant differences.

**Figure 5.**
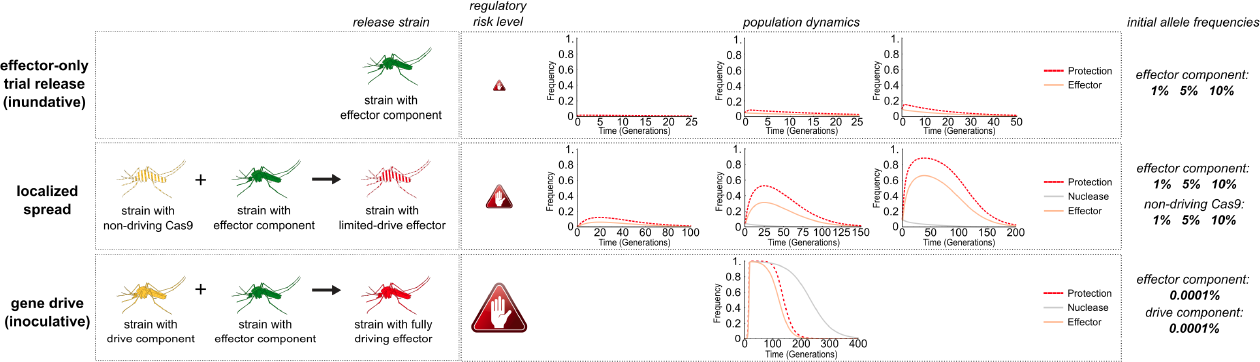
Evaluation of the allele frequency dynamics of limited release strategies. Release of strains carrying an IGD effector component only (upper panel) or of a limited-drive strain combining a non-driving Cas9 locus with an effector (middle panel) permit field testing of components in the absence of gene drive with the resulting changes in the target population dependent on the release size practically achievable. Release of the fully driving strain results in replacement of the wild-type population, even when released at low frequencies (lower panel).

## Discussion

The IGD paradigm reflects our view that a successful gene drive intervention will have to involve the use of multiple interacting traits rather than attempting to make singular monolithic constructs evolution-safe. It consequently should allow (informed by continuous monitoring) the ability to flexibly react to predictable as well as unexpected developments in a target vector or parasite population by adapting the release strategy in a context-dependent manner. Control would require the constant development and refinement of engineered genetic traits rather than the one-off or continued application of a static product.

Modelling of allele frequency dynamics suggests that for the parameters investigated, the IGD strategy could confer significant advantages over conventional replacement drive designs. By integrating components into endogenous loci, undesired repair outcomes following DNA cleavage tends to result in loss of function of the hijacked gene. Selection in turn reduces the rate at which such resistant alleles accrue. Whilst IGD does not prevent the onset of resistance, it could significantly extend the duration and level of protection available to a human population. This approach also resolves the longstanding issue of the arbitrary selection of a target sequence for population replacement where neutral sites show poor conservation whereas conserved sites are likely to be functionally constrained and thus costly to disrupt. The IGD strategy allows targeting a coding sequence of an endogenous gene while at the same time the insertion of IGD components is aimed to be neutral with respect to that gene, although it remains to be demonstrated experimentally if this is easily achievable. Notably, naturally occurring homing endonuclease genes, via their association with introns or inteins, propagate in a similar manner, i.e. by targeting highly conserved sites without disrupting their function.

There are several noteworthy assumptions made in the model. It is assumed following previous work (Beaghton et al. 2017a) that at baseline values, the cost of expressing the nuclease from the drive component (*s_n_*=0.05) is lower than the cost of expressing an effector gene from an effector component (*s_e2_*=*s_e3_*=0.1). This is justifiable when considering that germline genes tend to have lower expression levels than those present in the soma and that expression of antimalarial effectors needs to be sufficiently strong to ensure effective concentrations. It has been suggested that resistant alleles are susceptible to early stochastic loss (Beaghton, Beaghton, and Burt 2017) while we consider loss of gene function alleles to arise predominantly as a result of frameshift mutations, suggesting that some of the assumptions underpinning our deterministic model are conservative.

We have determined which parameters are key in influencing the duration of protection: the cost of expressing the nuclease (*s_n_*), the cost of disrupting the drive component locus (*s_d1_*), the cost of expressing the effector (*s_e_*), and the cost of disrupting its corresponding locus (*s_d2_*). Modeling shows changes in the cost of expressing the nuclease and the cost of disrupting the drive component locus are robustly tolerated. By contrast, the IGD strategy is sensitive to changes in the cost for disruption of the effector component locus, and the cost of expression of the effector. This is due to the efficiency with which the drive component, even if not present at high allele frequencies, is able to convert wild-type alleles into effector alleles. As such, significant penalties can be imposed upon the efficacy of the drive component without sizeable reduction in the generation of protection. These findings may be useful in the design of IGD traits, particularly with respect to the effector components. Minimizing the cost of effector expression and avoiding disruption of the hijacked gene are critical to the successful implementation of an IGD-based release. Previous work has highlighted the difficulty with which these parameters can be evaluated in a field setting (Z. N. Adelman and Tu 2016), however, the modularity of the IGD strategy may be an advantage in this respect.

The potential theoretical properties of IGDs should be seen in conjunction with the practical advantages of accelerating and simplifying the generation, testing, optimization, regulation and deployment of gene drives for population replacement. The IGD strategy offers an increased degree of versatility in testing and release of components. Modelling of effector-only and limited-drive releases suggests how IGD permits safe evaluation of effector components in the field, without need for a driving Cas9 source. This may alleviate concerns relating to the invasive nature of gene drives. From a logistical standpoint, the limited effect that such releases have on a localized test area facilitate the testing of multiple effectors in the same locale, once the presence of previously released transgenes have suitably diminished.

Our initial models included here must be extended to evaluate the potential of IGD in more realistic settings with full seasonality and spatial heterogeneity. Currently, spatial effects are not considered, and the simulated population is considered to be distributed homogeneously within a contained landscape. Previous work has been conducted to investigate the effects of spatial interactions upon the propagation of replacement drives (Eckhoff et al. 2017) and these models should be extended to IDGs. The inclusion of sex-specific parameters would allow the consideration of sex-specific effectors and associated fitness effects (Beaghton et al. 2017b). This is particularly relevant given that, in order to improve the efficiency and duration of the IGD strategy, it may prove useful to restrict effector activity to females and homing activity to males, reducing the net fitness cost on the population as a whole. Moreover, this approach may help to overcome issues associated with maternal deposition of Cas9 mRNA into the embryo (Champer et al. 2017a; A. M. Hammond et al. 2017b). We limited our analysis to one autonomous drive component and two non-autonomous effectors components. More powerful models could explore the dynamics of multiple drive and effector alleles and their interaction over more complex geographical scales and metapopulation structures. A set of separate hijacked host genes could be used in sequence to guarantee that the level of Cas9 in the population remains high even when at particular loci resistant alleles predominate in some areas or subpopulations. Equally, multiple effector strains could be generated expressing several effector molecules from different host genes and used to ensure that most individuals in each population carry transmission blocking effectors acting in various parasite-relevant tissues, even if resistant alleles are in circulation. Our simple 2-effector model already indicates that hedging homing over multiple loci in this manner could be a viable strategy to tackle the issues of resistance and standing variation in target populations, particularly as the use of multiple gRNAs at one site may only create a marginal difference to drive behaviour while significantly complicating construct design (Oberhofer, Ivy, and Hay 2018; Champer et al. 2018b). Finally, modelling could explore whether effector components could also be used in conjunction with Cas9-expressing suppressive strains to ensure a reduction in the vectorial capacity of mosquito populations that have been reduced in size but not eliminated. IGD population replacement could thus operate alongside a suppression program and would be a safeguard in the case of a population and transmission bounce following the intervention.

## Methods

### Discrete-generation population genetics model

The code for all models used in this study is available at https://github.com/genome-traffic/igd. The classical model for a drive+effector construct (see Model I, Beaghton et al. 2017) considers five different alleles: the wild-type allele (*w*), the complete drive construct which has both a functional nuclease and a functional effector (c=*n+e*), a nuclease only construct which has a functional nuclease but a defective effector (*n*), an effector only construct which has a functional effector but a defective nuclease (*e*), and functional resistant alleles (*r*), that are not recognized by the nuclease and have no functional nuclease or effector. Resistant alleles can either be pre-existing in the population or arise via NHEJ and MHMR repair pathways, as well as incomplete HDR. We define *d* as the transmission rate of drive, *u* as the parameter for resistance arising from end-joining repair, and *l_n_, l_e_*, and *l_ne_* as the probability of loss of function of effector, nuclease or both during HDR. Therefore, allele contributions from germline cleavage and homing from *w/c* are according to *w: c: n: e: r* in proportions (1 – *d*) (1 – *u*): *d* (1 – *l_n_ - l_e_* – *l_ne_*): *d* (*l_e_*): *d* (*l_n_*): (1- *d*) *u* + *l_ne_ d*, and in *w/n according to w: n: r* in proportions (1 – *d*) (1 - *u*): *d* (1 -*l_n_*): (1 – *d*) *u* + *l_n_ d.*

While the IGD model may involve multiple nuclease genes and effectors on many different loci, here we consider a simplified version with transgenes on either two or three independent loci *i*(with *i*=1,2,3). Locus 1 corresponds to the drive component, and loci 2 and 3 to effector components. At each locus *i*there are four possible alleles: a wild-type allele ( *w_i_*), a transgene *t_i_* (corresponding to either the nuclease gene as the transgene at the first locus, *t_1_* = *n*_1_, or to an effector component at the second or third locus, *t_2_ = e_2_* and *t_3_ = e_3_*), and two types of alleles at each locus that are resistant to the drive and do not have an intact nuclease or effector, *r_i_* and *m_i_*. The first type of resistant allele, *r_i_*, arises from incomplete homing or mutations that are in-frame and do not cause loss of the function of the host gene, and therefore are not considered to carry any fitness cost (similarly to the wild-type). The second type of resistant allele, *m_i_*, corresponds to a mutation that results in frameshift of the host gene, disrupting the endogenous locus. If the host gene is an essential gene, *m_i_* alleles are considered to convey lethality when homozygous. Resistance can be pre-existing or occur either by NHEJ or by incomplete HDR at either locus. We neglect resistant alleles created from spontaneous mutations, as rates are likely to be low compared to generation of resistance during homing.

An individual is considered to have IGD drive if it carries at least one functional drive component at locus 1 and at least one functional effector component at locus 2 (and at locus 3 if an additional effector is included as part of the strategy). This is a necessary condition for the effector component to home and propagate in the population at a super-Mendelian rate. For both models, we assume that the initial field population consists entirely of the wild-type allele, and there may be pre-existing resistance due to standing genetic variation at each locus (although the baseline pre-existing resistance is set to zero). Individuals homozygous for different IGD drive components are subsequently released as a relative proportion of the field population.

Cleavage and homing can occur only in the germline of genotypes with a wild-type and nuclease allele at locus 1 (*w*_1_/*n*_1_ at the first locus). Cleavage and homing at effector locus 2 (and locus 3 if an additional effector is included at another host gene) can occur only in those genotypes if there is at least one nuclease allele at locus 1 and a wild-type and effector allele at locus 2 (*w*_2_/*e*_2_ at locus 2). Transmission of the nuclease at locus 1 occurs with probability *d_1_*. Transmission of the effector transgene at locus 2 occurs with probability *d_2a_* when the drive component is heterozygous, and probability *d_2b_* when homozygous, and similarly for an effector at locus 3 if included (*d_3a_ d_3b_*). Resistance to the drive, sometimes accompanied by loss of gene function, is considered to occur during homing. We conservatively consider mutations to produce resistance (*r_i_*) in 1/3 of cases, and loss of gene function ( *m_i_*) in 2/3, here predominantly caused by frameshift mutations. Resistant alleles (*r_i_*) arise at loci 1,2, and 3 from incomplete HDR with probabilities *l_1_*, *l_2_* and *l_3_*, and by NHEJ with probabilities *u_1_*, *u_2_* and *u*_3_ respectively. Resistant alleles (*m_i_*) that cause loss of the endogenous gene function occur via incomplete HDR at loci 1,2, and 3 with probabilities *L_1_*, *L_2_* and *L_3_*, and by NHEJ with probabilities *U_1_*,*U_2_* and *U*_3_ respectively. Due to germline cleavage, homing, incomplete HDR and repair events, individuals that are heterozygous for the nuclease at locus 1, i.e. *w*_1_/*n*_1_, contribute alleles *w_1_: n_1_: r_1_: m_1_* in proportions (1 – *d_1_*) (1 – *u_1_* – *U_1_*): *d_1_* (1 – *l_1_ - L_1_*): (1 – *d_1_*) *u_1_* + *l_1_ d_1_*: (1 – *d_1_*) *U_1_* + *L_1_ d_1_.* Allele contributions from individuals with the wild-type and effector allele (*w*_2_/*e*_2_) at locus 2, if there are one or two nuclease alleles at locus 1, are according to *w*_2_: *e*_2_: *r*_2_: *m*_2_ in proportions (1 – *d_2k_*)(1- *u_2_* – *U_2_*): *d_2k_* (1 – *l_2_ - L_2_*): (1 – *d_2k_*) *u_2_* + *l_2_ d_2k_*: (1 – *d_2k_*) *U_2_* + *L_2_ d_2_* where *k* = a, b corresponds to locus 1 heterozygous or homozygous for the nuclease. If no nuclease allele is present at locus 1, gene transmission at locus 2 is Mendelian, as is inheritance in all other individuals. Similar expressions can be written for an additional effector at locus 3.

With four possible alleles at two independent loci, there are 16 gamete types and 100 diploid genotypes; for three independent loci, there are 64 gamete types and 1000 genotypes. The fitness of each genotype is relative to the wild-type homozygote ( *w_1_/w_1_; w_2_/w_2_; w_3_/w_3_)*, which has a fitness of one. Fitnesses are modelled using 10 parameters for transgenes at two loci and 14 parameters for three loci. We consider the following homozygous fitness costs: the cost of in-frame disruption (as caused by an intact transgene or resistant allele r at the host gene) at locus 1,2, and 3 (*s_d1_, s_d2_*, *s_d3_*), the cost of disruption (by *m**_i_*** mutations) that lead to loss of function at each locus (*s_m_*), the cost of expressing the nuclease from the drive component (*s_n_*), and the cost of expressing the effector from the effector components (*s_e2,_ s_e3_*). The corresponding dominance coefficients are *h_d1_*, *h_d2,_ h_d3,_ h_m_* and *h_n1_*, *h_e2_ h_e3_.* The range of fitness costs is from 0 (no cost) to 1 (lethal) and dominance coefficients range from 0 (completely recessive) to 1 (completely dominant). The fitness of each genotype is derived as the product of costs at each locus associated with site disruption, number of nuclease components, and number of effector components. For example, for a two-loci model (drive at locus 1 and effector at locus 2), the fitness of a genotype that is heterozygous for the transgene at both loci (*w_1_/n_1_; w_2_/e_2_*) is given by (1 – *h_d1_ s_d1_*) (1 – *h_d2_ s_d2_*) (1 – *h_n_ s_n_*)( 1 – *h_e2_ s_e2_*) to reflect costs of host gene disruption by transgenes (for baseline parameters, this cost is set to zero) at both loci as well as the cost of expressing the nuclease (at locus 1) and the effector (locus 2).

Allele frequencies and genotype abundances are modelling using deterministic discrete-generation recursion equations. We assume a one life stage model (adults) with a field population composed of equal numbers of male and females with the same genetic and fitness parameters, such that allelic and genotypic frequencies are equal between them. Mating is random, with unsuccessful mating events not considered. We assume the population to be sufficiently large to ignore stochastic effects. The system of equations is solved numerically using Wolfram Mathematica (Wolfram Research, Inc., Mathematica, Version 11.3, Champaign, IL (2018)).

As in Beaghton et al. (2017), the effect of the IGD strategy on transmission of disease is dependent on the frequency of each genotype in the population, and the reduction in vector competence when one or more effector components is present. For the 2-loci model, the reduction is denoted by *h_rc1_r_c_* if the effector component is present at locus 2 in one copy (heterozygous) and by *r_c_* if two copies (homozygous) are present. For the 3-loci model, we assume that the reduction in vector competence depends on the total number of effector alleles, giving *h_rci_ r_c_* for *i* =1, 2 or 3 effector alleles in total over loci 2 and 3, and *r_c_* for 4 alleles in total (i.e., homozygous for the effector element in both loci). Values of *r_c_* range from 0 (no effect) to 1 (total transmission blockage), and the dominance coefficient for refractoriness *h_rc1_* (and *h_rc2_ and h_rc3_* for the 3-loci model) ranges from 0 (completely recessive) to 1 (completely dominant). We quantify the effect in terms of the reduction in vectorial capacity at time *t* as 1 – *V_C_*[t], where *V_C_*[t] is the vectorial capacity. *V_C_*[t] is calculated as the sum over the genotype frequencies multiplied by their individual vector competence. For the 2-locus model, this yields:

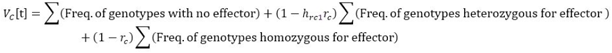

For the 3-locus model:

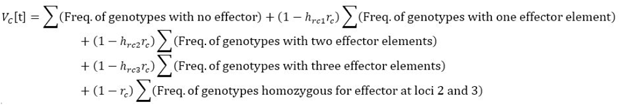

## Acknowledgements

This study was funded by the Bill & Melinda Gates Foundation grant OPP1158151 awarded to NW & GC. Funded by the European Research Council under the European Union's Seventh Framework Programme ERC grant no. 335724 awarded to N.W. A.N. is supported by the Biotechnology and Biological Sciences Research (BBSRC) Council-funded Doctoral Training Partnership award 1655064.

---

**Supplementary Figure 1.**
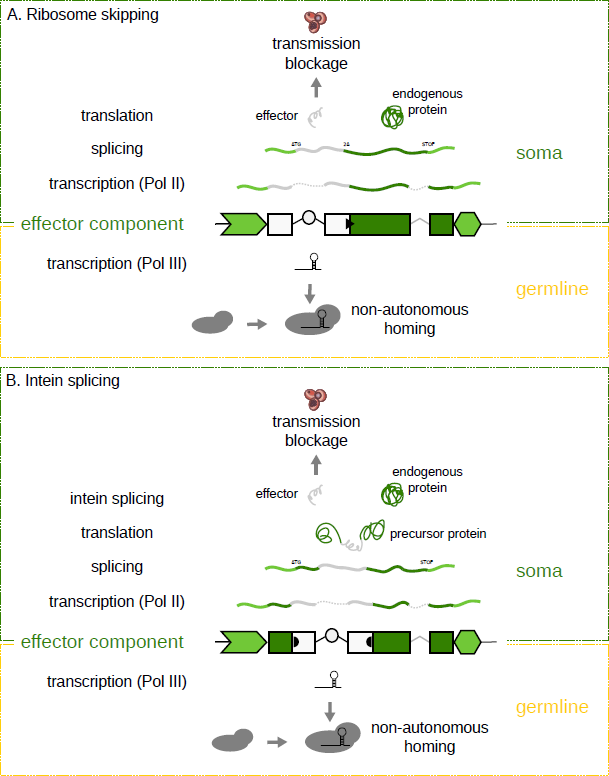
Schematic showing the expression of the endogenous and introduced genetic elements of an effector component allele at a hijacked gene locus. Expression of the gRNA occurs in the germline (yellow) whereas expression of the hijacked gene occurs in an infection-relevant somatic tissue (green). While expressing a straight fusion of endogenous protein (green) and the effector protein (grey) is possible the generation of separate polypeptides by using either (A) the 2A ribosome-skipping signal (black triangle) or (B) the use of intein sequences (black semicircle) are possible avenues to avoid interfering with the function of the hijacked gene.

**Supplementary Figure 2.**
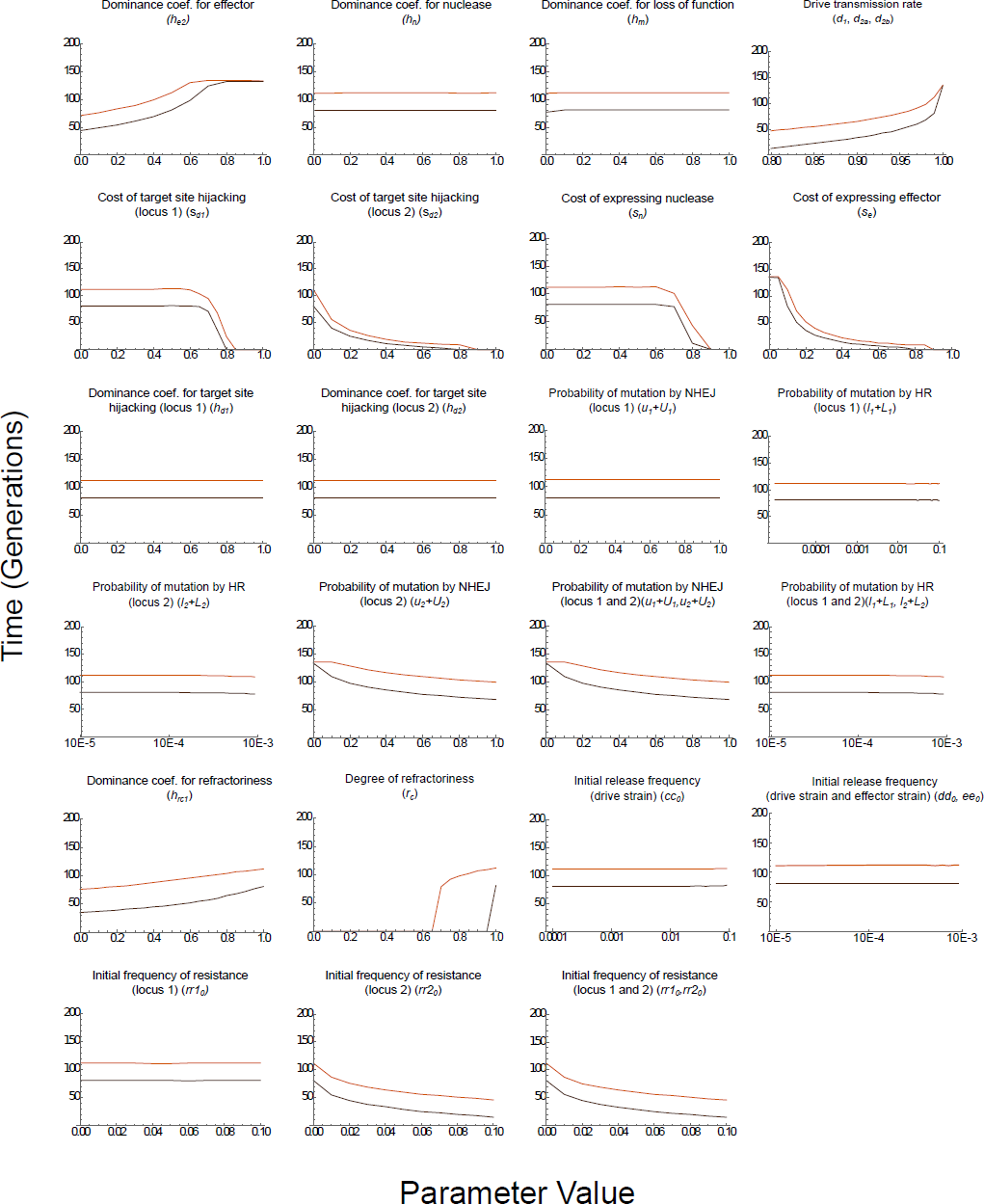
Duration of 67% and 95% protection (top and bottom lines in each graph) as each of the underlying parameters is varied, holding all others at their baseline values. Note that the drive parameters (d10). For variation of 1), with the ratio of resistance vs loss of gene function held at ½.

**Supplementary Figure 3.**
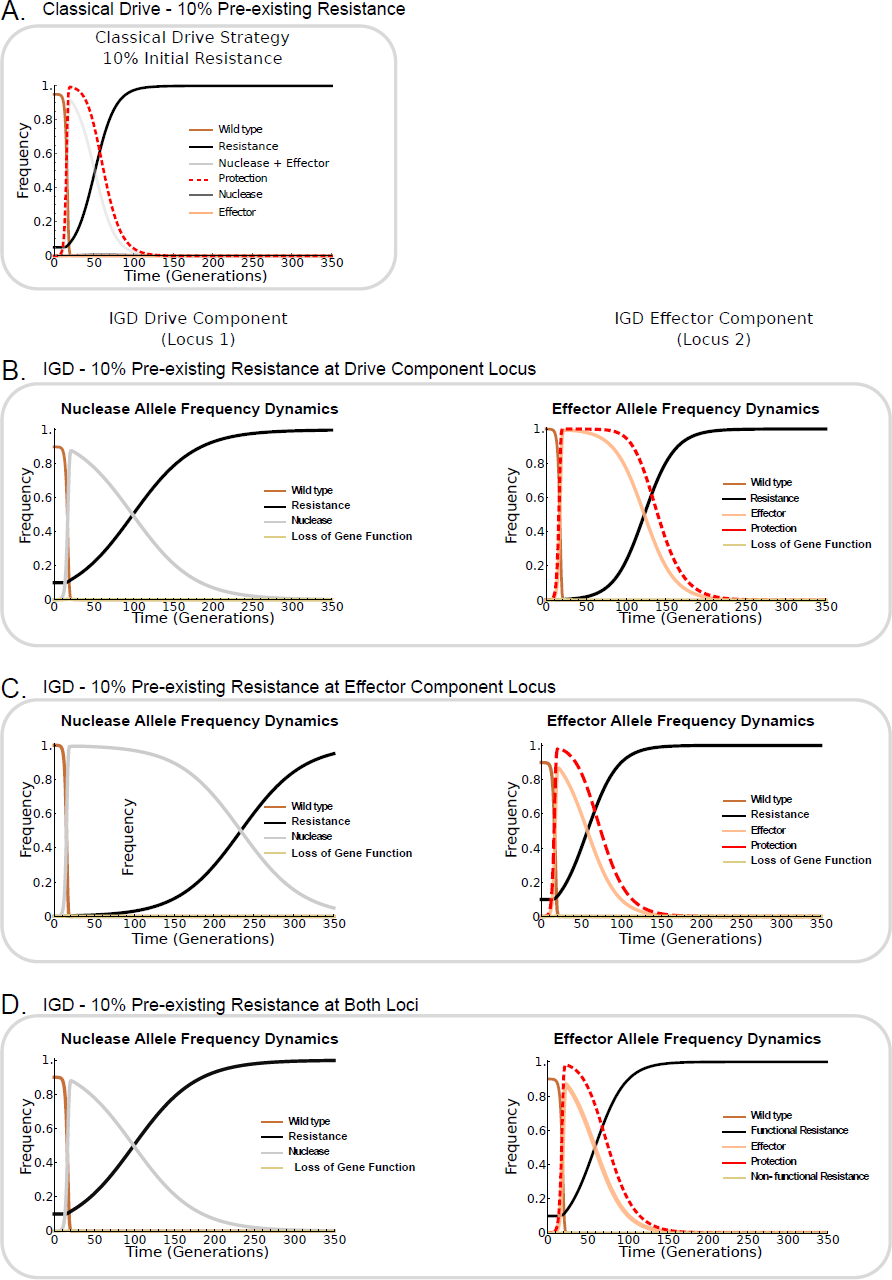
Overview of allele frequency dynamics when releasing both classical drive (A) and IGD strains into field populations with 10% pre-existing resistance. For IGD, resistance is modelled at the drive component locus (B), the effector component locus (C), and at both loci (D).

